# Kappa opioid receptors control a stress-sensitive brain circuit and drive cocaine seeking

**DOI:** 10.1101/2025.06.23.661181

**Authors:** Valentina Martinez Damonte, Lydia G. Bailey, Amit Thakar, Joanna Stralka, Travis E. Brown, Julie A. Kauer

**Affiliations:** Department of Psychiatry and Behavioral Sciences, Nancy Pritzker Laboratory, Stanford University, Stanford, CA 94305, US; Department of Integrative Physiology and Neuroscience, Washington State University, Pullman, WA 99163, US

**Keywords:** Ventral tegmental area, nucleus accumbens, optogenetics, self-administration, LTP, plasticity

## Abstract

Stress is a potent trigger for drug-seeking behaviors in both rodents and humans with a history of substance use. Kappa opioid receptors (kORs) play a critical role in mediating stress responses. Our previous studies in the ventral tegmental area (VTA) demonstrated that acute stress activates kORs to block long-term potentiation at GABA_A_ synapses on dopamine neurons (LTP_GABA_) and triggers stress-induced reinstatement of cocaine seeking. Here we identify the specific GABAergic afferents affected by stress, the precise localization of kORs within the VTA, and show that VTA kOR activation is sufficient to drive reinstatement.

We optogenetically activated specific GABAergic afferents and found that nucleus accumbens (NAc)-to-VTA, but not lateral hypothalamus (LH)-to-VTA projections, exhibit stress-sensitive LTP_GABA_. Using a conditional knock-out approach, we found that selectively deleting kORs from NAc neurons but not from dopamine cells prevents stress-induced block of LTP_GABA_. Selectively activating dynorphin-containing NAc neurons with an excitatory DREADD mimics acute stress, preventing LTP_GABA_ at VTA synapses. We furthermore demonstrated that without acute stress, microinjection of a selective kOR agonist directly into the VTA facilitates cocaine reinstatement without similarly affecting sucrose-motivated responding, demonstrating the critical role of kORs in stress-induced cocaine reinstatement.

Our results show that kORs on GABAergic NAc nerve terminals in the VTA underlie loss of LTP_GABA_ that may drive stress-induced addiction-related behaviors. Our work highlights the importance of inhibitory inputs for controlling dopamine neuron excitability in the context of addiction and contributes to defining the circuit involved in stress-induced drug reinstatement.

## INTRODUCTION

Stressful events cause profound changes in reward-related behaviors. Exposure to brief stress triggers robust relapse to drug-seeking in animals when cocaine self-administration was previously extinguished and can also trigger relapse in humans (Mantsch et al., 2016). This connection between stress and substance use disorder has sparked interest in the mesolimbic reward circuitry as a potential site for stress-induced dysregulation. Although typically linked to reward and reinforcement, ventral tegmental area (VTA) dopamine neurons also show increased activity in response to acute and chronic stress (Holly and Miczek, 2016). VTA dopamine neuron excitability is tightly controlled by inhibitory synapses from multiple local and distant afferent sources (Kaufling et al., 2010; Watabe-Uchida et al., 2012; Beier et al., 2015; Beier et al., 2019), and accordingly, the removal of inhibition onto dopamine neurons enhances dopaminergic activity (Johnson & North, 1992; Paladini & Tepper, 1999; Tan et al., 2012; van Zessen et al., 2012; Xin et al., 2016; Simmons et al., 2017). Moreover, synaptic plasticity is selectively expressed at subsets of inhibitory synapses on dopamine neurons (Simmons et al., 2017, Polter et al., 2018, St Laurent and Kauer, 2019; Martinez Damonte et al., 2023), which participate in functionally diverse circuits (Sesack & Grace, 2010; Beier et al., 2015; Beier et al., 2019; Heymann et al., 2020). This heterogeneity indicates that specific synapses on distinct subpopulations of dopamine neurons will likely be selectively and differentially modified by drugs or stress.

We and others have shown that GABAergic synapses and plasticity of these synapses in the VTA contribute to the development of drug reinforcement and stress-induced relapse to drug seeking (Laviolette and van der Kooy, 2001; Laviolette et al., 2004; Nugent et al., 2007; Niehaus et al., 2010; Tan et al., 2010; Graziane et al., 2013; Beier et al., 2017; Polter et al., 2014, 2017). Either brief cold-water swim stress or exposure to addictive drugs persistently blocks a form of synaptic plasticity at GABAergic synapses on VTA dopamine neurons (LTP_GABA_) (Nugent et al., 2007; Niehaus et al., 2010; Graziane et al., 2013, Polter et al., 2014, 2017). The stress-induced block of LTP_GABA_ is triggered by the activation of VTA kappa opioid receptors (kORs), which remain persistently activated. Even days after an acute stressor, interrupting kOR signaling both rescues LTP_GABA_ in brain slice recordings and prevents reinstatement of cocaine-seeking *in vivo* (Polter et al., 2014, 2017).

Kappa OR agonists alone can induce dysphoria in humans (Pfeiffer et al., 1986), aversion and depressive-like behaviors in rodents (Ebner et al., 2010; Chefer et al., 2013; Ehrich et al., 2015), and reinstate cocaine self-administration in non-human primates (Valdez et al., 2007). Further, manipulation of kORs influences drug-seeking behaviors both during and after drug use cessation (Bruchas et al., 2010). Systemic kOR activation reinstates cocaine conditioned place preference (CPP), enhances nicotine CPP, and reinstates seeking for nicotine, ethanol, and amphetamines (Schenk & Partridge, 2001; Redila & Chavkin, 2008; Smith et al., 2012; Funk et al., 2014; Grella et al., 2014; Harshberger et al., 2016). Kappa OR antagonists are therefore currently in clinical trials for treating substance use disorders (Lobe et al., 2025). Defining the involvement of specific subpopulations of kORs and the circuits they modulate may ultimately lead to more specific treatments with fewer off-target effects.

Here we identified a stress-sensitive circuit that may drive stress-induced addiction-related behaviors. Kappa ORs located on nucleus accumbens (NAc) neurons that project to the VTA —but not on dopamine neurons—become activated by stress to block LTP_GABA_ at these synapses. Moreover, chemogenetic activation of dynorphin-expressing neurons in the NAc mimics the effect of stress on LTP_GABA_. Finally, we report that simply activating kORs in the VTA is sufficient to trigger reinstatement of cocaine-seeking without acute stress, highlighting the potential of VTA kOR modulation for controlling addictive behaviors.

## METHODS

### Animals

All procedures were carried out in accordance with the guidelines of the National Institutes of Health for Animal Care and Use, and they were also approved by the Administrative Panel on Laboratory Animal Care from Stanford University.

Electrophysiological experiments in this study used several mouse lines: DAT^IRESCre^ knock-in (referred to as DatCre, Jackson Laboratory, stock number: 006660, strain name: B6.SJL-Slc6a3^tm1.1(cre)Bkmn^/J), Vgat-IRES-cre knock-in (Jackson Laboratory, stock number: 028862, strain name: B6J.129S6(FVB)-Slc32a1^tm2(cre)Lowl/^MwarJ), Pdyn-IRES-Cre (Jackson Laboratory, stock number: 027958, strain name: B6;129S-Pdyn^tm1.1(cre)Mjkr^/LowlJ), Pitx3-GFP (Jackson Laboratory, stock number: 41479-JAX, strain name: B6.129P2-Pitx3^tm1Mli^/Mmjax), (Zhao et al., 2004), kOR^fl/fl^ (Jackson Laboratory, stock number: 030076, strain name: B6;129-Oprk1^tm2.1Kff^/J), here referred to as kOR ^-/-^ (Jackson Laboratory, stock number: 035045, strain name: STOCK Oprk1tm1.1(cre)Sros/J) and C57BL/6 male and female mice bred in-house. The mouse strain used for this research project, B6.129P2-*Pitx3^tm1Mli^*/Mmjax, RRID:MMRRC_041479-JAX, was originally obtained from the Mutant Mouse Resource and Research Center (MMRRC) at The Jackson Laboratory, an NIH-funded strain repository, and was donated to the MMRRC by Meng Li, BMed, PhD, Cardiff School of Biosciences (Zhao et al., 2004). Additionally, some experiments required mice that lack kOR in dopamine cells, e.g. mice that were heterozygous for the loxP-flanked kOR allele and heterozygous for the cre transgene in dopamine neurons. kOR ^-/-^ mice, which are homozygous for loxP-flanked kOR, were mated to DatCre mice. Approximately 50% of the offspring was heterozygous for the loxP allele and heterozygous for the cre transgene (F1). These mice were mated back to the homozygous loxP-flanked mice. Approximately 25% of the progeny from this mating (F2) was homozygous for the loxP-flanked allele and heterozygous for the cre transgene, which constituted the experimental mice. Mice were maintained on a 12-h light/dark cycle and provided food and water *ad libitum*. Both male and female mice were used for experiments.

Behavioral experiments utilized adult male Sprague-Dawley rats (Envigo) weighing 230-350g at the time of surgery. Rats were maintained on a reverse 12-h light/dark cycle (lights off at 08:00), in a temperature and humidity-controlled colony room, with *ad libitum* access to food and water until behavioral testing began, at which point animals were restricted to 20g of chow/day. All training occurred during the animals’ dark phase (08:00-20:00).

### Acute stress protocol

Acute stress was delivered using a modified Porsolt forced swim task. Mice were placed in a 2 L plastic beaker containing approximately 1 L of cold water (4-6 °C) for 5 min under continuous monitoring. Following the swim task, animals were dried off, wrapped in a dry cloth for 5 min, placed singly in a warmed cage that was heated underneath by a heating pad for 2 h and then mice were returned to the home cage. Midbrain slices were prepared 24 h after stress exposure.

Footshock stress was delivered in operant chambers equipped for mice (MedAssociates). Mice were given a mild (0.21mA) footshock (Soria et al., 2008) once every minute for 5 minutes, and then individually housed in a standard home cage for 24 h until euthanasia for electrophysiology. Control animals were placed in the same operant chamber for 5 minutes but received no shock.

### Preparation of brain slices

Brain slices were prepared from mice deeply anesthetized using ketamine. An ice-cold oxygenated solution was delivered via cardiac perfusion (in mM): 110 choline chloride; 25 glucose; 25 NaHCO_3_; 7 MgCl_2_; 11.6 sodium ascorbate; 3.1 sodium pyruvate; 2.5 KCl; 1.25 NaH_2_PO_4_ and 0.5 CaCl_2_. Following perfusion, the brain was rapidly dissected and horizontal slices (220 μm) through the VTA were prepared using a Leica Microsystems vibratome. Slices were then transferred to a holding chamber where they were submerged in artificial cerebrospinal fluid containing (aCSF, in mM): 126 NaCl, 21.4 NaHCO_3_, 2.5 KCl, 1.2 NaH_2_PO_4_, 2.4 CaCl_2_, 1.2 MgSO_4_, 11.1 glucose and 0.4 ascorbic acid, saturated with 95%O_2_/5% CO_2_ (pH 7.4) at 37°C for 10 minutes. Slices were then held at room temperature until transferred to the recording chamber.

### Electrophysiology

Midbrain slices were submerged in a recording chamber and continuously perfused at 1.5–2 ml/min with aCSF at 28–32°C containing: 6,7-dinitroquinoxaline-2,3-dione (DNQX; Tocris; 10μM), (2R)-amino-5-phosphonopentanoate (APV; Tocris; 100 µM) and strychnine (Sigma-Aldrich; 1μM) to block AMPA-, NMDA- and glycine receptors, respectively. Patch pipettes (2-5 MΩ) were filled with (Sigma-Aldrich; in mM): 125 KCl, 2.8 NaCl, 2 MgCl2, 2 ATP-Na+, 0.3 GTP, 0.6 EGTA, and 10 HEPES (pH=7.25-7.28; 265-280 mOsm; junction potential at 30°C 3.4 mV). Whole-cell patch-clamp recordings were made from neurons visually identified in the lateral VTA.

Dopamine neurons were identified by the presence of a large I_h_-current (>25 pA) during a voltage step from −50 mV to −100 mV (Margolis et al., 2003). For consistency with our previous work we followed this criterion, aware that a subset of the neurons recorded from and reported here are possibly non-dopaminergic neurons (Margolis et al., 2006). Additionally, in Figures 1, 2 and 5 we used a dopamine neuron reporter mouse, PITX3-GFP (Zhao et al., 2004). Dopamine cells were voltage-clamped at -70mV and cell input resistance and series resistance were monitored throughout the experiment using a -10 mV hyperpolarizing step from -70 mV for 100 ms. Cells were discarded if these values changed by more than 15% during the experiment. IPSCs were amplified using an AM Systems amplifier, low-pass filtered at 3 kHz and digitally sampled at 30 kHz using an analog-to-digital interface (Digidata; Molecular Devices).

**Figure 1.**
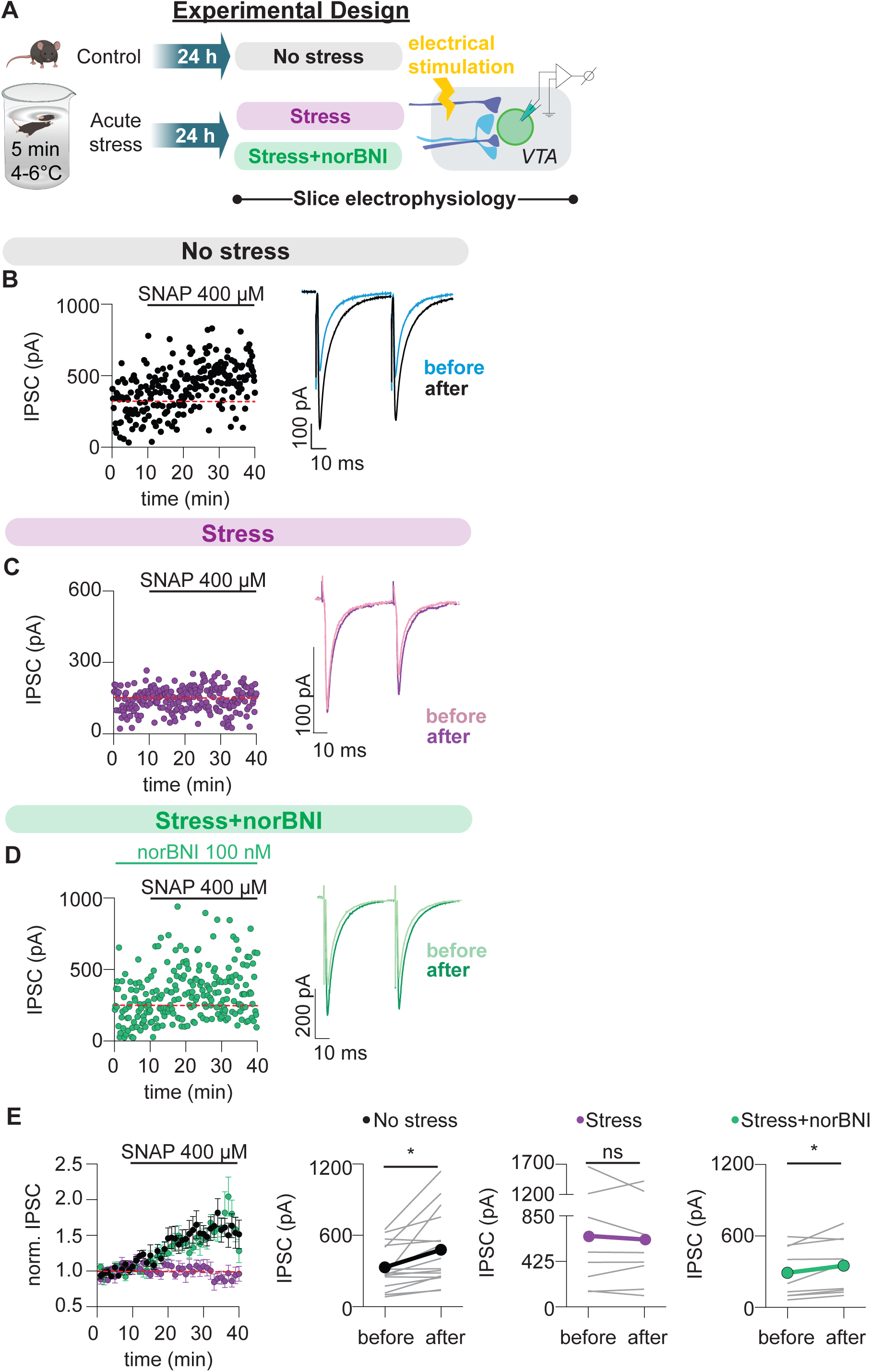
GABAergic IPSCs in mice display nitric oxide-induced long term potentiation (LTP_GABA_). (A) Experimental design. Brain slices were prepared 24 hr after acute stress or no stress; recordings from dopamine neurons were carried out with or without norBNI. (B, C, D) Representative time courses and example IPSCs from electrophysiological recordings in slices from (B) control mice, (C) mice stressed 24 hr previously, and (D) mice stressed 24 hr previously with bath application of nor-BNI to the slice. All IPSCs in this and other figures were evoked with a rostrally-placed electrical stimulating electrode. (E) Time courses of averaged IPSC amplitudes (left) and IPSC amplitudes before and after SNAP application (right) in neurons in unstressed (n = 16 cells/10 mice, 12 cells from male mice, 4 from female mice, Paired t test, p = .007), stressed (n = 8 cells/8 mice, 6 cells from male mice, 2 from female mice, Paired t test, p = .61) or stressed + norBNI conditions (n = 7 cells/7 mice, 5 cells from male mice, 2 from female mice, Paired t test, p = .04).

**Figure 2.**
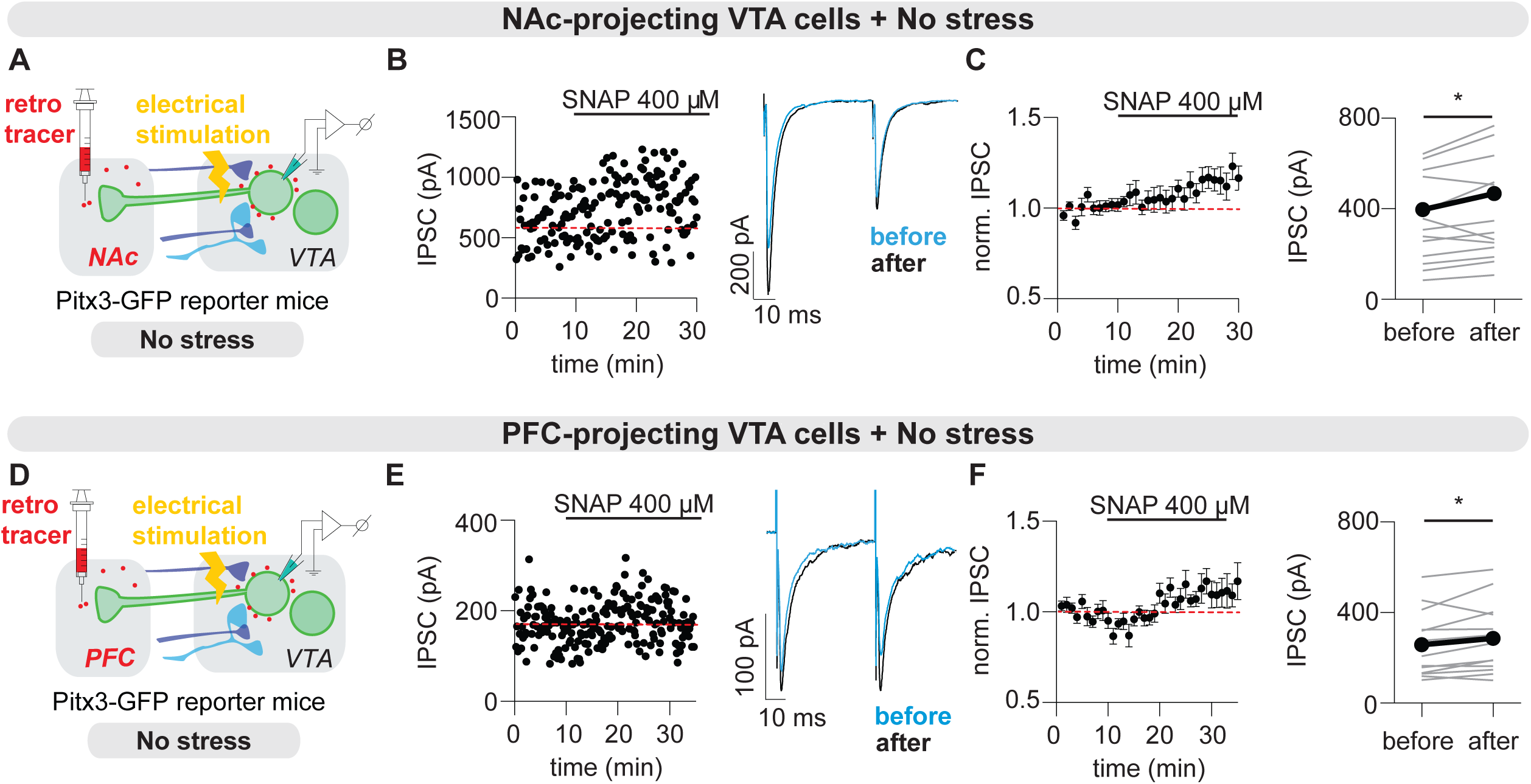
NAc- and PFC-projecting VTA dopamine neurons receive synapses that display LTP_GABA_. (A) Experimental design for (B) and (C). Neurons from NAc-projecting DA neurons were identified in slices with retrobeads. (B) Representative time course and example IPSCs from NAc-projecting DA neurons before and after SNAP application. (C) Time courses of averaged IPSC amplitudes (left) and IPSC amplitudes before and after SNAP application (right) (NAc-projecting, n = 14 cells/12 mice, 10 cells from males, 4 from females, Paired t test, p = .04). (D) Experimental design for (E) and (F). Neurons from PFC-projecting DA neurons were identified in slices with retrobeads. (E) Representative time course and example IPSCs from PFC-projecting DA neurons before and after SNAP application. (F) Time courses of averaged IPSC amplitudes (left) and IPSC amplitudes before and after SNAP application (right) (PFC-projecting, n = 13 cells/10 mice, 5 cells from males, 8 from females, Paired t test, p = .04.).

### Stimulation protocols

For electrical stimulation, a bipolar stainless-steel stimulating electrode was placed rostral to the VTA 200-500 μm from the recorded cell. GABA_A_ receptor-mediated IPSCs were evoked at 0.1 Hz using 100 μsec current pulses.

Channelrhodopsin-induced synaptic currents were evoked at 0.033 Hz using 0.1- to 5-ms full-field light pulses from an LED (Mightex), filtered and controlled by driver (ThorLabs) and reflected through a 40x water immersion lens. Pairs or trains of light-evoked IPSCs (oIPSCs) were separated by 30s to avoid desensitization of the opsin. For all stimulus frequencies, intensity remained constant throughout the experiment. Long-term potentiation was elicited by bath application of S-nitroso-N-acetylpenicillamine (SNAP, Tocris; 400 μM) after obtaining a stable 10 min baseline (Nugent et al., 2007).

### Stereotaxic injections

Stereotaxic surgeries were performed on male and female mice between postnatal days 25-35. For experiments involving selective optogenetic activation of NAc or LH inputs to the VTA, Vgat-IRES-cre::PITX3-GFP mice were deeply anesthetized with isofluorane and 200-300 nL of AAV-DJ-hSyn-hChR2(H134R)-eYFP were bilaterally injected into the NAc (AP:+1.55, ML: ±1.15, DV:-4.60) or the LH (AP:-1.55, ML: ±1.10, DV:-5.20). Instead, for chemogenetic experiments, we used Pdyn-IRES-Cre mice and AAV-DJ-EF1a-DIO-hM3D(Gq)-mCherry. Horizontal slices containing the VTA were prepared 4-7 weeks after surgery. For experiments involving retrograde labeling of dopamine cells, 200-300 nL of Retrobeads^TM^ (Lumafluor) or DiI were injected into the NAc or into the PFC (AP:+2.00, ML: ±0.20, DV:-3.00). Slices from injection regions were also prepared to confirm the injection locations; slices from mice with mistargeted injection sites of virus or of retrobeads were discarded.

### *In situ* hybridization

Mice brains were dissected and immersed in pre-chilled isopentane for freezing and were then embedded in OCT within cryomolds. Horizontal brain sections (20 µm) were obtained using a cryostat (Leica Biosystems), mounted onto glass slides, and dried for 1 hr within the cryostat before being transferred to −80°C for long-term storage. Fluorescence *in situ* hybridization was performed using the RNAscope Multiplex Fluorescent Reagent Kit v2 (Advanced Cell Diagnostics (ACD), Cat. No. 323136), specifically following the *Fresh Frozen Sample Preparation and Pretreatment* protocol provided by the kit. The following probes from ACD were used: Mm-Slc6a3 (Cat. 315441); Mm-Slc32a1-C2 (Cat. 319191-C2), and Mm-Oprk1-O2-C3I (Cat. 1573161-C3). Images were captured using a Keyence BZ-X800 fluorescent microscope (Keyence) using the 40x objective.

### Jugular Catheter Surgery

Jugular catheterization surgery was conducted as described previously (McFarland & Kalivas 2001). Rats were anesthetized with a zyket cocktail (90 mg/kg ketamine; 10 mg/kg xylazine) given intraperitoneally prior to implanting a chronic indwelling intravenous catheter.

Catheters were constructed with silastic tubing (10 cm, outer diameter 0.047 in, inner diameter 0.025 in) connected to a back-mount cannula pedestal. The pedestal was constructed in-house using a 10mm fully threaded nylon stud and 25 mm long 22-gauge hypodermic tubing, which protruded 5 mm from the top of the nylon stud, and the bottom was bent to 90 degrees. The pedestal and tubing were encased within an acrylic base with polypropylene mesh. A small ball of silicone sealant was placed at 2.8 mm from the end of the silastic tubing. Tubing was inserted into the jugular vein up to the silicone sealant ball and sutured securely to the underlying muscle tissue.

Immediately after jugular catheter surgery, catheters were flushed with 0.1 mL cefazolin (100 mg/mL in sterile saline) and 0.05 mL Taurolidine-Citrate Solution (TCS; BrainTree Scientific, Braintree, MA).

Rats were then fixed in a stereotaxic frame, and in the event of a toe-pinch reflex, maintained on a low dose (0.5-1%) of isoflurane through a nosecone for the remainder of the surgery. Cannulas were implanted bilaterally into the VTA at 10° angles (AP:-5.2, ML: ±1.8, DV: -7.5; Wang et al., 2012). Cannulas were cemented into place using dental cement, and sutures placed. Rats were given 5 mg/kg carprofen (NSAID), 0.65 mg/kg Ethiqua (opioid), 1 mL of sterile saline, and local anesthetic on incision sites. Animals were kept on a heating pad until anesthetics wore off. For one week following surgery, animals were assessed daily to ensure health, including weight checks and saline injections to aid recovery when needed.

One week following surgery, catheter patency was assessed using a 0.1 mL injection of propofol (10 mg/mL), followed by 0.1 mL of cefazolin and 0.05 mL TCS. Cefazolin and TCS injections then occurred daily immediately following self-administration sessions to reduce the chance of infection and to maintain catheter patency.

### Drugs

Drugs were made up as stock solutions and diluted to appropriate concentrations in aCSF (*in vitro* use) or saline (*in vivo* use). S-nitroso-N-adenylpenacillamine (SNAP; Tocris; 400 μM) was made up in DMSO and bath applied to elicit LTP. Where indicated, norbinaltorphimine (norBNI, 100 nM) was bath applied to slices at least 10 min prior to induction of LTP_GABA_. For chemogenetic experiments in Figure 6, clozapine dihydrochloride (CLZ; Hello Bio) at a dose of 0.1 mg/kg in sterile saline OR saline was delivered via intraperitoneal (i.p.) injections 24 h before electrophysiological recordings. For the behavioral experiment, ±-U-50488 hydrochloride (Tocris, Bristol, UK) was dissolved in sterile saline, filtered, and administered at 0.3 μg/0.5 μL/hemisphere.

### Self-Administration

Experiments were conducted in operant-conditioning chambers enclosed in sound-minimizing cabinets (Med-Associates, Inc.). Each chamber contained an active and inactive lever and corresponding cue lights, a house light, and a food dispenser.

Each box was equipped with a variable speed pump, which was controlled by the self-administration chamber.

Throughout self-administration, animals were maintained on 20g of chow/day, given immediately following their session in the self-administration chambers. Training began 7-9 days following surgery. All training occurred during the animals’ dark phase (08:00-20:00). Animals were placed in the same chamber at the same time daily for 2 h.

#### Training

Animals underwent a minimum of 6 consecutive days of shaping to encourage active lever responding, which consisted of alternating cardboard and cardboard + sucrose on the active lever. All training was on a fixed ratio 1 (FR1) schedule of reinforcement. An active lever response resulted in cocaine infusion (0.2 mg/kg/infusion over 2.6 seconds) and illumination of the cue light over the correct lever, and a tone cue for 5 seconds. After a correct lever response, a timeout period of 30 seconds occurred during which responses on either lever were counted, but no rewards or cues were given. Inactive lever responses were counted, but no consequences were given. Once animals received more than 10 cocaine infusions for two consecutive days using these incentives, cardboard and sucrose were removed, and animals were allowed to continue responding for cocaine. Animals were required to have 10 consecutive days of more than 10 cocaine rewards to transition to the extinction phase.

Animals that did not have functioning catheters underwent the same training, except that they received a sucrose pellet from the food hopper instead of an infusion of cocaine. All protocols and criteria were otherwise kept the same for both groups.

#### Extinction

During extinction training sessions, lever pressing resulted in no infusions of cocaine or activation of the cue light or sound. The house light was still active in all sessions. Animals underwent a minimum of 7 days of extinction training and continued until reaching two consecutive days of fewer than 10 active lever presses. On day 4 of extinction, dummy cannulas were removed, and injector needles were placed in the cannulas for 1 minute. On day 5 of extinction, animals received a sham saline injection through their cannulas prior to their extinction session. These procedures were to minimize the stress of handling and injection to reduce the influence of these procedures on testing days.

#### Reinstatement Testing

Once animals had two consecutive days of fewer than 10 active lever responses, reinstatement was tested. Animals received an intra-VTA injection of either U50488H (0.3μg/hemisphere in 0.5 μL sterile saline; Pham et al., 2021; Zhang & Kelley, 1997; Castro & Berridge, 2014) or sterile saline (0.5 μL/hemisphere) over the course of 1 minute and given 1 minute for absorption prior to removing injector needles. 10 minutes following injection, animals were given a 2-hour reinstatement test, during which active lever responses resulted in the cue light and tone, but no infusion of cocaine.

### Cannula Placement Verification

For the behavioral experiment, following the conclusion of reinstatement testing animals received an intra-VTA injection of 2 μL of 2.5% dextran tetramethylrhodamine to visualize placement. Animals were then euthanized, and brains were fixed with 4% paraformaldehyde overnight and switched to 20% sucrose in phosphate buffered saline for another 24 h. Brains were sliced at 40-micron thickness, and placements were visualized using a fluorescence microscope to ensure accurate placement of cannulas.

### Electrophysiological data analysis

IPSC peak amplitudes were measured, and responses were normalized by taking the mean of the 10-min baseline responses and dividing the rest of the responses by this mean. These normalized values were then used for average plots. For these plots, all cells were time aligned at the beginning of the 10-min baseline and averaged over the entire period.

The expression of LTP was determined comparing averaged IPSC amplitudes for 5 min just before LTP induction with averaged IPSC amplitudes during the 15-20 min period after manipulation using a paired Wilcoxon test with a significance level of p˂0.05.

### Experimental Design and Statistical Analysis

Sample sizes were determined from our previous experiments and from related literature for electrophysiology experiments and behavior. All data are presented as mean ± standard error. Differences were deemed significant with p < .05. Data were first tested using the Shapiro-Wilk test for assessing normality. Data that passed this test were tested using a parametric test, while data that did not display a normal distribution were tested using a nonparametric test for significance.

## RESULTS

We used whole-cell patch clamp recordings from mouse VTA dopamine neurons to test whether electrically evoked inhibitory postsynaptic currents (IPSCs) were potentiated by the nitric oxide donor, SNAP. In these and all subsequent experiments, inhibitory synaptic currents were isolated using AMPA and glycine receptor antagonists. As we have shown before in rats (Nugent et al., 2007; Graziane et al., 2013; Polter et al., 2014), we found that under control conditions (no stress) in mice GABAergic IPSCs evoked with a rostrally-placed electrical stimulating electrode also display nitric oxide-induced long-term potentiation (LTP_GABA_) (Figure 1A-B,E). As in rats, this form of plasticity in VTA slices is entirely blocked 24 h after a 5 min cold-water swim stress exposure (Figure 1C,E), and is rescued with bath application of the kOR inverse agonist, norbinaltorphimine (norBNI) (Figure 1D,E). Further studies determined that a footshock stressor exhibits a similar LTP_GABA_ block; 24 h after mice were given a mild (0.21mA) footshock stressor, VTA neurons showed a lack of SNAP induced LTP (data not shown; main effect of treatment F(1, 12) = 9.4, p=0.0098), suggesting that footshock stress exhibits a similar effect on VTA LTP_GABA_ as forced swim stress.

Over the last few decades, it has become clear that VTA dopamine neurons are a highly heterogeneous population, with distinct projection targets and participating in distinct circuits that are differentially modulated (Lammel et al., 2014). To test whether dopamine neurons projecting to either NAc or prefrontal cortex (PFC) participate in subcircuits sensitive to acute stress, we injected a retrograde tracer into each of these two main target regions of VTA dopamine neurons to label dopamine neurons projecting to each region, and later tested for nitric-oxide induced LTP in the two dopamine neuron subpopulations (Figure 2). We found that electrically evoked IPSCs in both NAc- and PFC-projecting VTA dopamine neurons can display LTP_GABA_. The LTP elicited in PFC-projecting cells was smaller than what we usually observe in unidentified VTA dopamine neurons, perhaps because our sampled PFC-projecting neurons were restricted to those that exhibit I_h_ and often were located in lateral VTA, whereas most PFC projecting VTA dopamine neurons exhibit little I_h_ and are found in the medial VTA (Lammel et al., 2014); it is possible that a subset of dopamine neurons we sampled is not representative of all PFC-projecting neurons.

We reported previously that a kOR agonist delivered *in vitro* prevented LTP_GABA_ (Graziane et al., 2013), and interrupting kOR signaling in slices from previously stressed animals reinstates the ability of an NO donor to produce LTP_GABA_ (Figure 1D; Polter 2014). These data indicate that during the exposure to acute stress, dynorphin is released within the VTA, activating local kORs in the region. Here we confirmed that kORs are required for LTP_GABA_ using global kOR^-/-^ mice that lack expression of kOR in every cell type (Figure 3). To test the efficacy of kOR deletion, we performed in situ hybridization in horizontal brain slices from wild type and kOR^-/-^ mice. While wild type mice displayed kOR expression in the VTA, this transcript is absent in kOR^-/-^ mice (Figure 3B and Supplemental Figure 1). Consistent with our previous evidence, in slices from these animals, exposure to stress did not prevent LTP_GABA_ at 24 h (Figure 3C-D).

**Figure 3.**
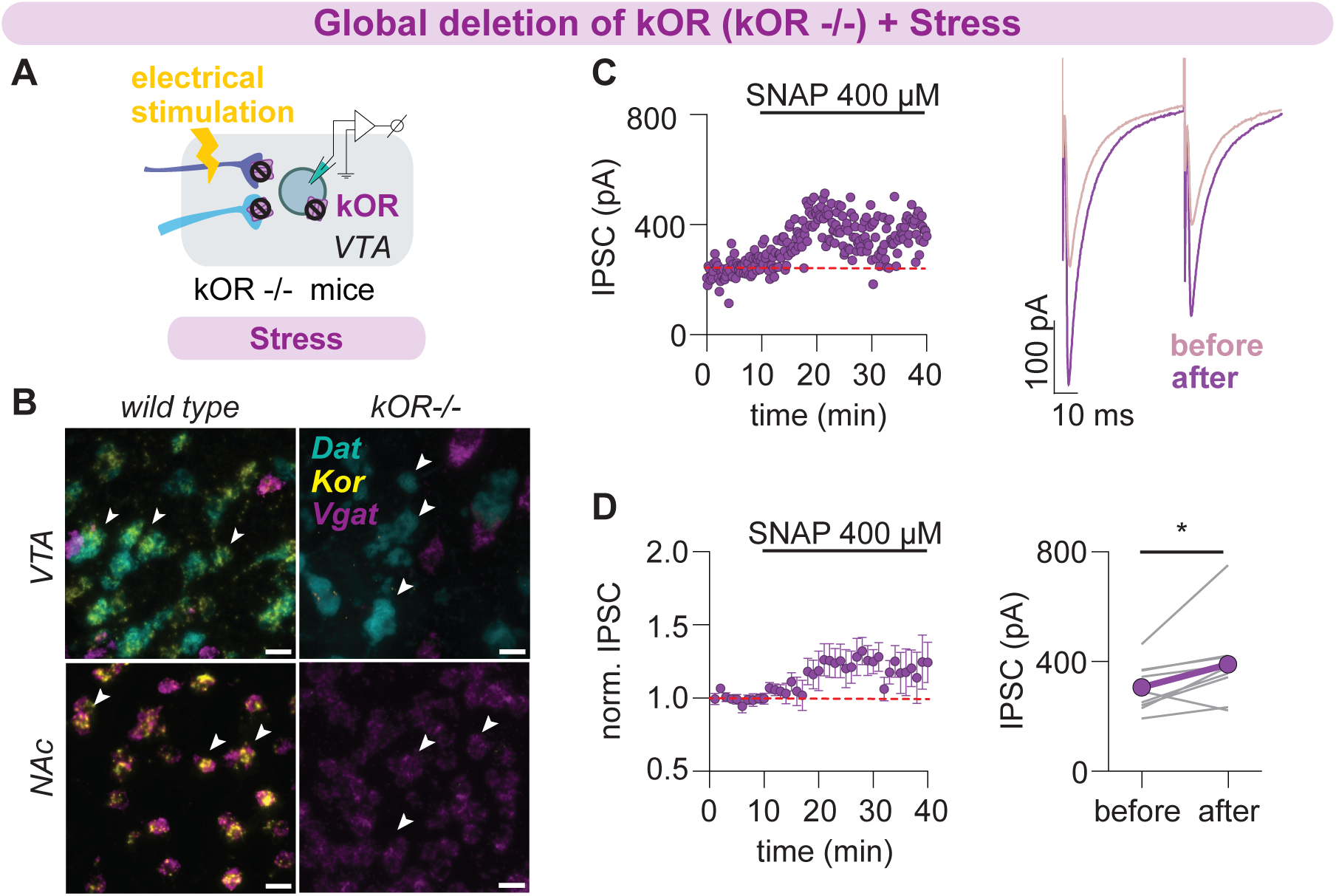
Global deletion of kORs prevents stress-induced block of LTP_GABA_. (A) Experimental design using slices from global knockout mice (kOR ^-/-^). (B) Representative confocal images showing horizontal VTA and NAc sections from wild type and kOR ^-/-^ mice validating global deletion of kORs. Scale bar 30 *μ*m. (C) Representative time course and example IPSCs before and after SNAP application in a brain slice from a kOR^-/-^ mouse. (D) Time courses of averaged IPSC amplitudes (left) and IPSC amplitudes before and after SNAP application (right) in recordings from neurons from kOR ^-/-^ mice stressed 24 hr previously (n = 9 cells/7 mice, 6 cells from male mice, 3 from female mice, Wilcoxon matched-pairs signed rank test, p = 0.04.)

We predicted that the block of LTP_GABA_ by acute stress would be blocked if kORs in the relevant cell type were deleted. Kappa ORs are expressed on dopamine neurons in the VTA as well as on multiple afferent fibers (Margolis et al., 2003; Abraham et al., 2018). To localize which of these kOR populations is responsible for the stress-induced block of LTP_GABA_, we next used an intersectional genetic approach to delete the kORs from specific candidate cell groups. To remove kOR selectively from dopaminergic neurons, we crossed mice expressing cre recombinase in dopamine neurons (DAT-cre) with mice in which the kOR gene is flanked by loxP sites (kOR^fl/fl^) (Figure 4A-B). We reasoned that if the kORs on dopamine cells are responsible for stress-induced loss of LTP_GABA_, in these mice (kOR^-/-DA^) LTP_GABA_ should become impervious to acute swim stress. Instead, conditional deletion of kOR from dopamine neurons did not prevent block of LTP_GABA_ after acute stress, demonstrating that the kORs present on dopaminergic neurons do not mediate stress-triggered loss of LTP_GABA_ (Figure 4C-E).

**Figure 4.**
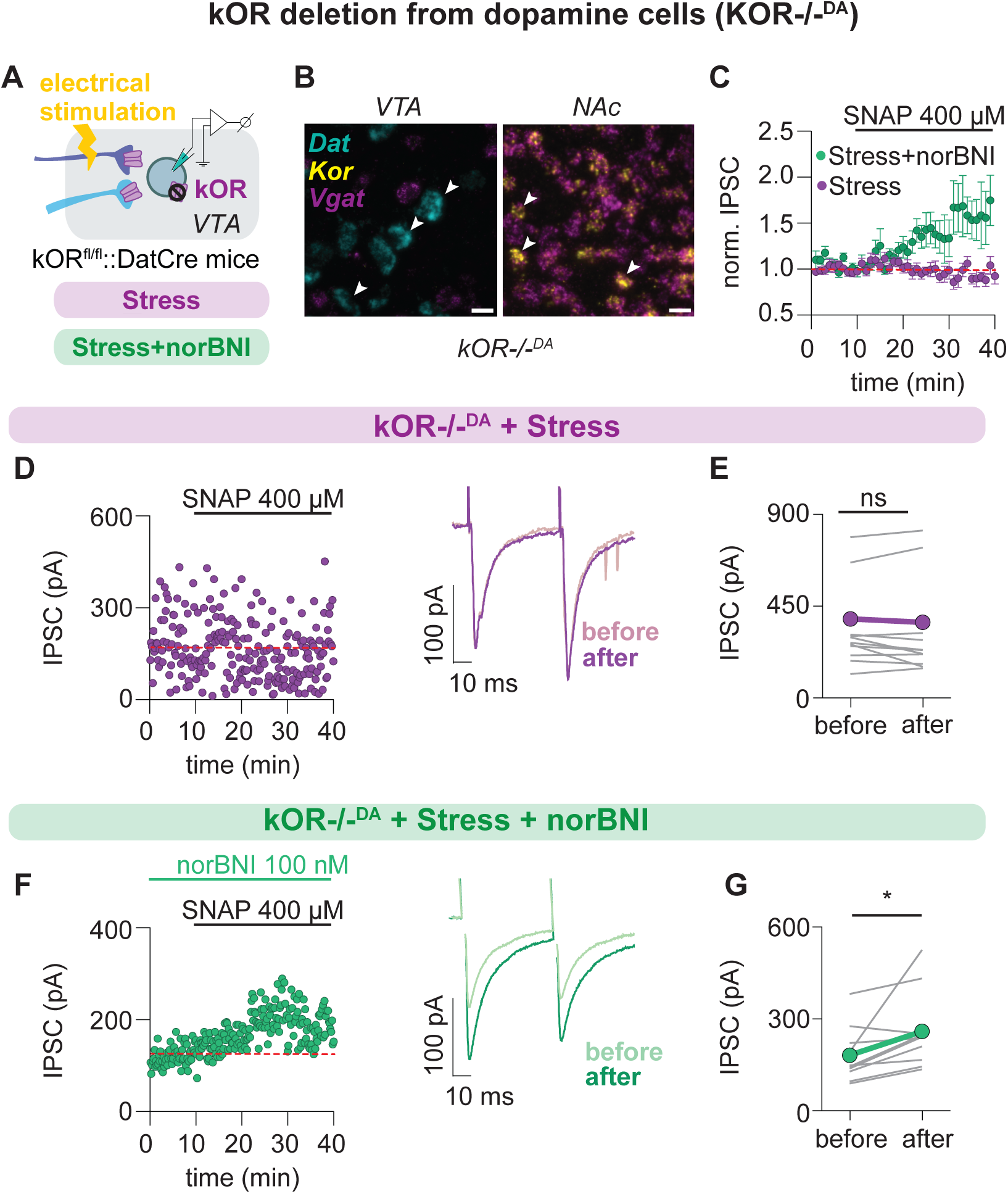
Deleting kORs from dopamine neurons does not prevent stress-induced block of LTP_GABA_. (A) Experimental design using mice that lack kOR in dopamine neurons (kOR^-/-DA^). (B) Representative confocal images showing horizontal VTA and NAc sections from kOR^-/-DA^ mice validating selective deletion of kORs from dopamine neurons. Scale bar 30 *μ*m. (C) Time courses of averaged IPSC amplitudes before and after SNAP application 24hr after stress for (D,E) and (F,G) experiments. (D) Representative time course, example IPSCs and (E) IPSC amplitudes before and after SNAP application in recordings from neurons from kOR^-/-DA^ mice 24 hr after stress. (n = 12 cells/11 mice, 5 cells from male mice, 6 from female mice, Wilcoxon matched-pairs signed rank test, p = 0.34). (F) Representative time course, example IPSCs and (G) IPSC amplitudes before and after SNAP application in recordings in the presence of norBNI from neurons from stressed kOR^-/-DA^ mice (n = 10 cells/9 mice, 5 cells from male mice, 5 from female mice, Paired t test, p = .01).

Additionally, norBNI bath application to slices lacking kORs in dopamine neurons rescued LTP_GABA_ as in wild type mice (Figure 4C, F-G), again indicating the presence of a separate kOR population in the VTA responsible for the loss of LTP_GABA_ after acute stress.

We next tested the possibility that the relevant kORs are located on the presynaptic GABAergic terminals that potentiate during LTP_GABA_. It is well established that different GABAergic afferents to the VTA exhibit distinct forms of synaptic plasticity (Dacher and Nugent, 2011; Simmons et al., 2017; Polter et al., 2018; St. Laurent and Kauer, 2019; St. Laurent et al., 2020; Martinez Damonte et al., 2023). We tested whether nitric-oxide-evoked LTP_GABA_ is present in two major GABAergic projections to the VTA involved in motivated behavior, the lateral hypothalamus (LH) and the nucleus accumbens (NAc). We used AAVs to express channelrhodopsin in neurons in either the LH or the NAc, and then in VTA slices we tested for SNAP-induced LTP of optically-evoked IPSCs (oIPSCs) in dopamine neurons (Figure 5A-B). Optically evoked IPSCs (oIPSCs) evoked by brief light pulses to ChR2-expressing LH-to-VTA afferents did not potentiate in response to bath-applied SNAP (Figure 5C). This result indicates that LH afferents are unlikely to contribute to the LTP_GABA_ seen in electrically evoked IPSCs, nor to its loss after acute stress. In contrast, oIPSCs evoked by light in ChR2-expressing NAc-to-VTA afferents were nearly always potentiated after exposure to SNAP, supporting the hypothesis that these synapses contribute to electrically-evoked LTP_GABA_ (Figure 5D-F), consistent with similar previous findings (Simmons et al., 2017).

Importantly, LTP_GABA_ at these synapses was sensitive to stress; 24 h after a 5 min acute swim stress LTP was not observed in the same optically stimulated NAc-to-VTA synapses (Figure 5G-I). Together, these data support the hypothesis that synapses on dopamine neurons from GABAergic afferents from the NAc are normally capable of exhibiting nitric oxide-triggered LTP, but that a single acute stress experience is sufficient to block that LTP.

These data suggested that the relevant kORs for stress-induced loss of LTP_GABA_ may be located on NAc terminals in the VTA. To test this, we co-injected a virus to drive expression of cre recombinase along with a second virus coding for cre-dependent channelrhodopsin (Figure 5J) into the NAc of kOR^fl/fl^ mice to selectively delete kORs from infected NAc neurons (kOR-/-^NAc^) and enabling us to activate these neurons with light. Simply deleting kORs from nerve terminals of NAc neurons entirely prevented stress from blocking LTP_GABA_ (Figure 5K-L). This result demonstrates a key role for NAc projections to VTA dopamine neurons in dynorphin sensitivity during acute stress.

**Figure 5.**
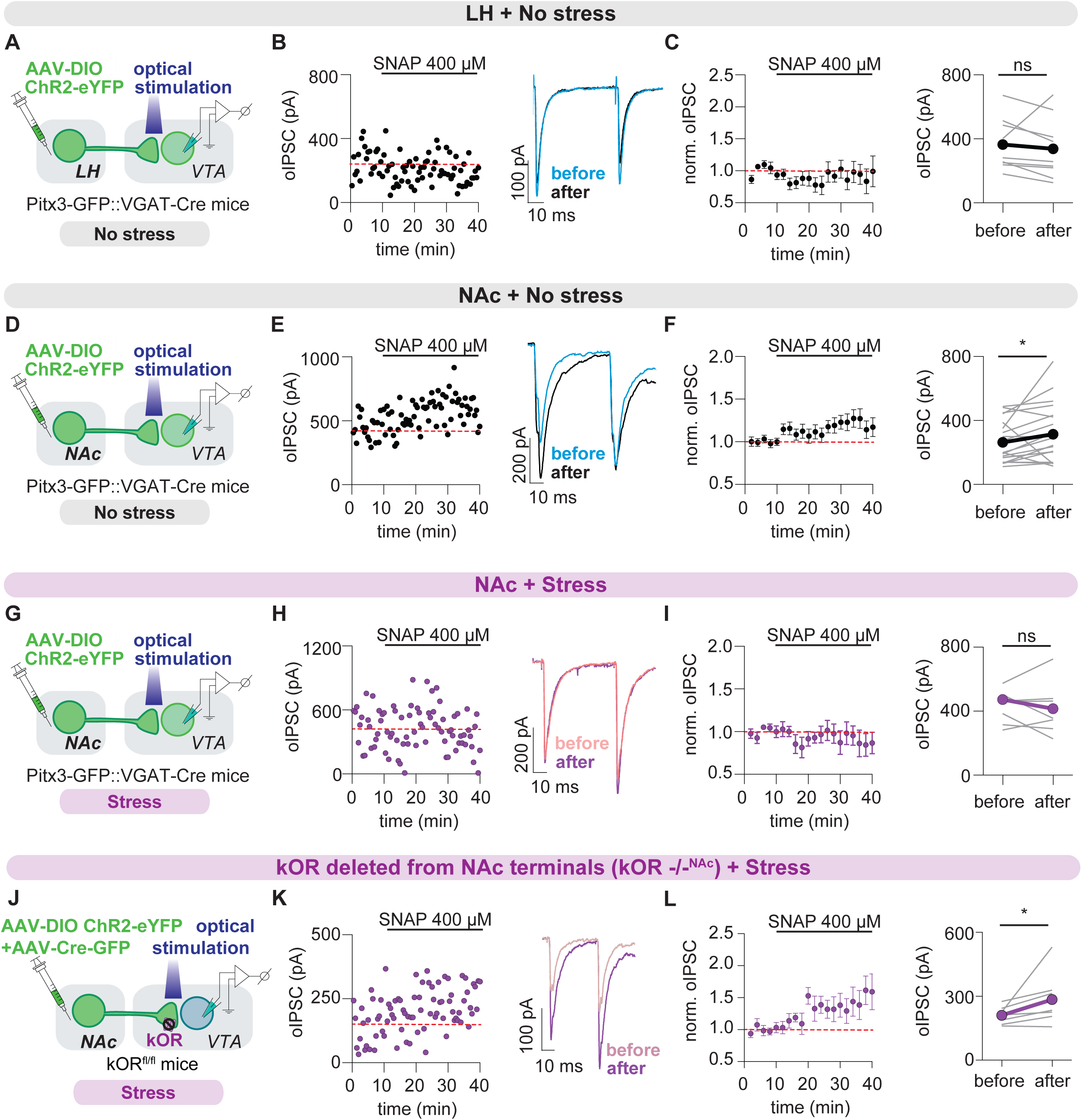
NAc-to-VTA but not LH-to-VTA GABAergic afferents display stress sensitive LTP_GABA_, which is lost upon deleting kOR from NAc neurons. (A) Experimental design for (B) and (C). (B) Representative time course and example LH-oIPSCs, and (C) time course of averaged LH-oIPSC amplitudes and LH-oIPSC amplitudes before and after SNAP application from control unstressed mice (n = 9 cells/7 mice, 4 cells from male mice, 5 from female mice, Paired t test, p = .51). (D) Experimental design for (E) and (F). (E) Representative time course, example NAc-oIPSCs, and (F) time course of averaged NAc-oIPSC amplitudes and NAc-oIPSC amplitudes before and after SNAP application from unstressed control mice (n = 20 cells/17 mice, 12 cells from male mice, 8 from female mice, Wilcoxon matched-paired signed rank test, p = .03). (G) Experimental design for (H) and (I). (H) Representative time course, example NAc-oIPSCs, and (I) time course of averaged NAc-oIPSC amplitudes and NAc-oIPSC amplitudes before and after SNAP application from mice stressed 24 hr previously (n =8 cells/6 mice, 5 cells from male mice, 3 from female mice, Paired t test, p = .35). (J) Experimental design for (K) and (L) using mice in which we optogenetically drove GABAergic NAc terminals from which kORs were deleted (kOR ^-/-NAc^). (K) Representative time course and example oIPSCs before and after SNAP application from stressed mice lacking kORs in NAc terminals in the VTA. (L) Time course of averaged oIPSC amplitudes and oIPSC amplitudes before and after SNAP application from mice stressed 24 hr previously (n = 7 cells/6 mice, 6 cells from male mice, 1 from a female mouse, Wilcoxon matched-pairs signed rank test, p = .03).

Our data indicate that dynorphin is likely released within the VTA following acute stress, and that dynorphin activates kORs on GABAergic afferents originating in the NAc. This activation prevents LTP_GABA_, for many days (Polter et al., 2014, 2017). Which dynorphin-containing afferents release dynorphin during acute stress? The VTA receives dynorphin afferents from several brain regions including the NAc D1R medium spiny neurons (MSNs), most of which co-express dynorphin (Fallon et al., 1985; Hara et al., 2006). We therefore next asked whether activation of dynorphin-containing neurons in the NAc is sufficient to mimic the effect of stress on LTP_GABA_. We injected the NAc of mice expressing cre recombinase in prodynorphin neurons (pdyn-cre mice, Krashes et al., 2014) with a virus that encodes for the activating DREADD, hM3Dq (Figure 6A). We predicted that upon activation of the DREADD, NAc dynorphin-containing neurons would be excited, releasing dynorphin from their nerve terminals in the VTA. After 3 weeks of DREADD expression, we injected the mice with either saline or clozapine to test whether activating these neurons would mimic acute swim stress. Remarkably, 24 hours later we found that while slices from saline-injected mice exhibited normal LTP_GABA_, injecting CNO to selectively activate dynorphin-containing neurons in the NAc *in vivo* mimicked acute swim stress, blocking LTP_GABA_ in slices (Figure 6B-F). This result suggests that not only do VTA projecting NAc neurons undergo stress-sensitive LTP_GABA_, but they also may be responsible for the release of dynorphin during acute stress.

**Figure 6.**
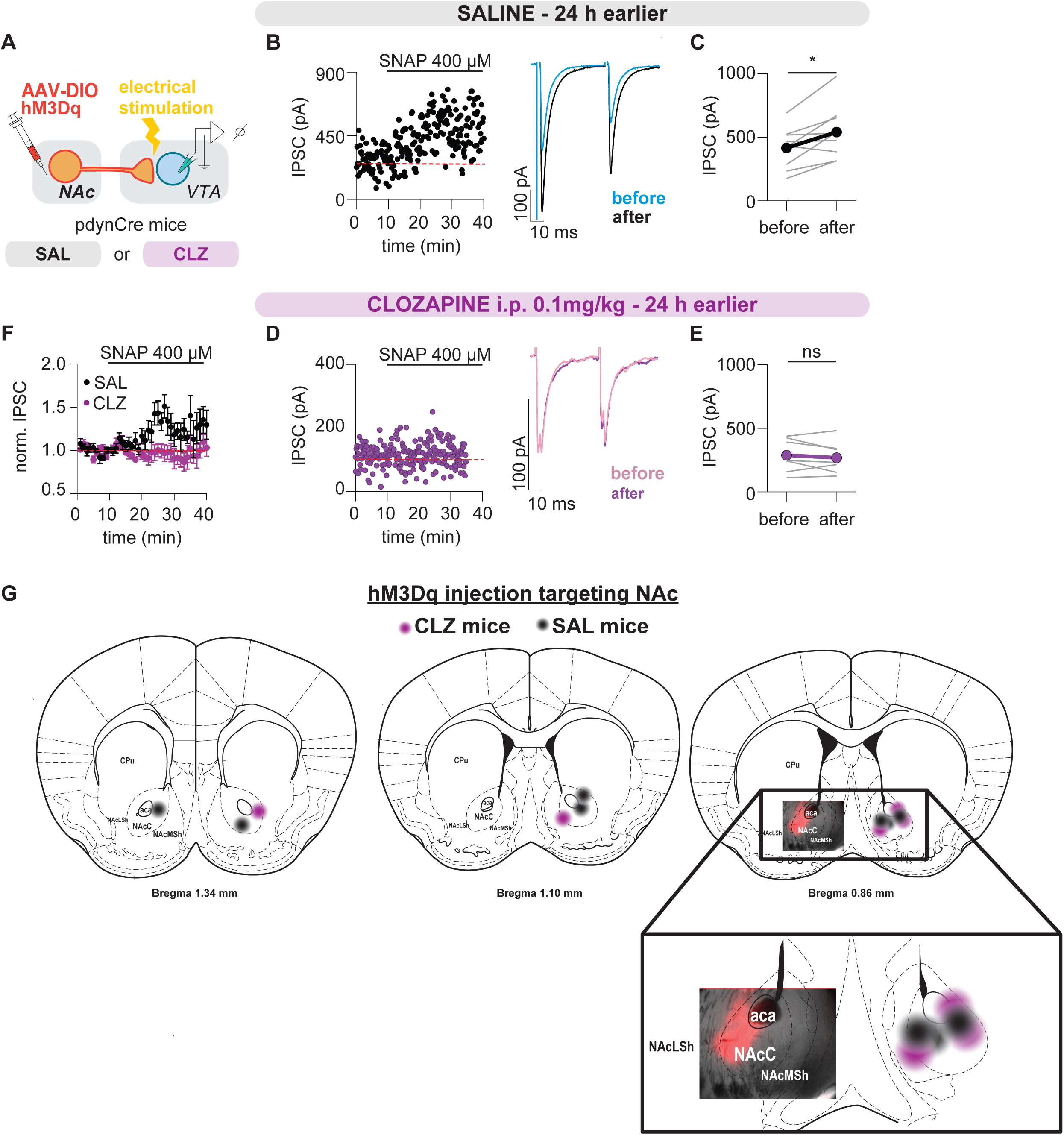
Chemogenetically activating dynorphin-containing neurons in the NAc mimics stress-induced block of LTP_GABA_. (A) Experimental design. Chemogenetic activation of pdyn neurons in the NAc. (B) Representative time course and example IPSC amplitudes and (C) IPSC amplitudes before and after SNAP application in slices from unstressed mice that received saline 24 h before slice preparation (n = 9 cells/7 mice, 4 cells from males, 5 from females, Paired t test, p = .01) . (D) Representative time course and example IPSC amplitudes and (E) IPSC amplitudes before and after SNAP application in slices from unstressed mice that received clozapine 24 h before slice preparation (n = 7 cells/5 mice, 4 cells from males, 3 from females, Paired t test, p = .44) . (F) Time courses of averaged IPSC amplitudes for (C) and (E). (G) Schemes illustrating hM3Dq injection targeting NAc from all animals.

Our previous work showed that stress-induced reinstatement of cocaine-seeking correlates well with the stress-induced kOR-dependent loss of LTP_GABA_. To determine the behavioral implications of our electrophysiological findings, we investigated whether pharmacological activation of VTA kORs with the kOR agonist, U50488H, would induce reinstatement of cocaine-seeking in rats previously trained to self-administer cocaine (Figure 7A). Prior to U50488H microinjection, no significant differences were observed between groups 1 and 2 in active or inactive lever responses, or cocaine infusions (Figures 7C & 7D, all p>0.44). Furthermore, there were no significant differences in lever responses on the first or last day of extinction (p>0.68) or the number of days to reach extinction criteria (p=0.79, data not shown) between groups. As expected, both groups showed a decrease in the number of active lever responses on the last day of extinction compared to the first day of extinction (Figure 7E: 2-way RM ANOVA; main effect of day: F_1, 14_=47.24, p<0.0001). After extinction, however, bilateral intra-VTA microinjection of U50488H increased cocaine-seeking compared to saline-microinjected controls (Figure 7F: Mann-Whitney U=8, p=0.01). In addition to total lever responses, the temporal pattern of responding differed over the 2-hour reinstatement test with U50488H microinjection eliciting a more robust and persistent cocaine-seeking behavior throughout the session (Figure 7G: 2-way RM ANOVA; main effect of treatment: F_1, 14_=4.86, p=0.04; main effect of time: F_11, 154_=2.87, p=0.002). These data show that activation of kORs within the VTA reinstates cocaine-seeking behavior.

**Figure 7.**
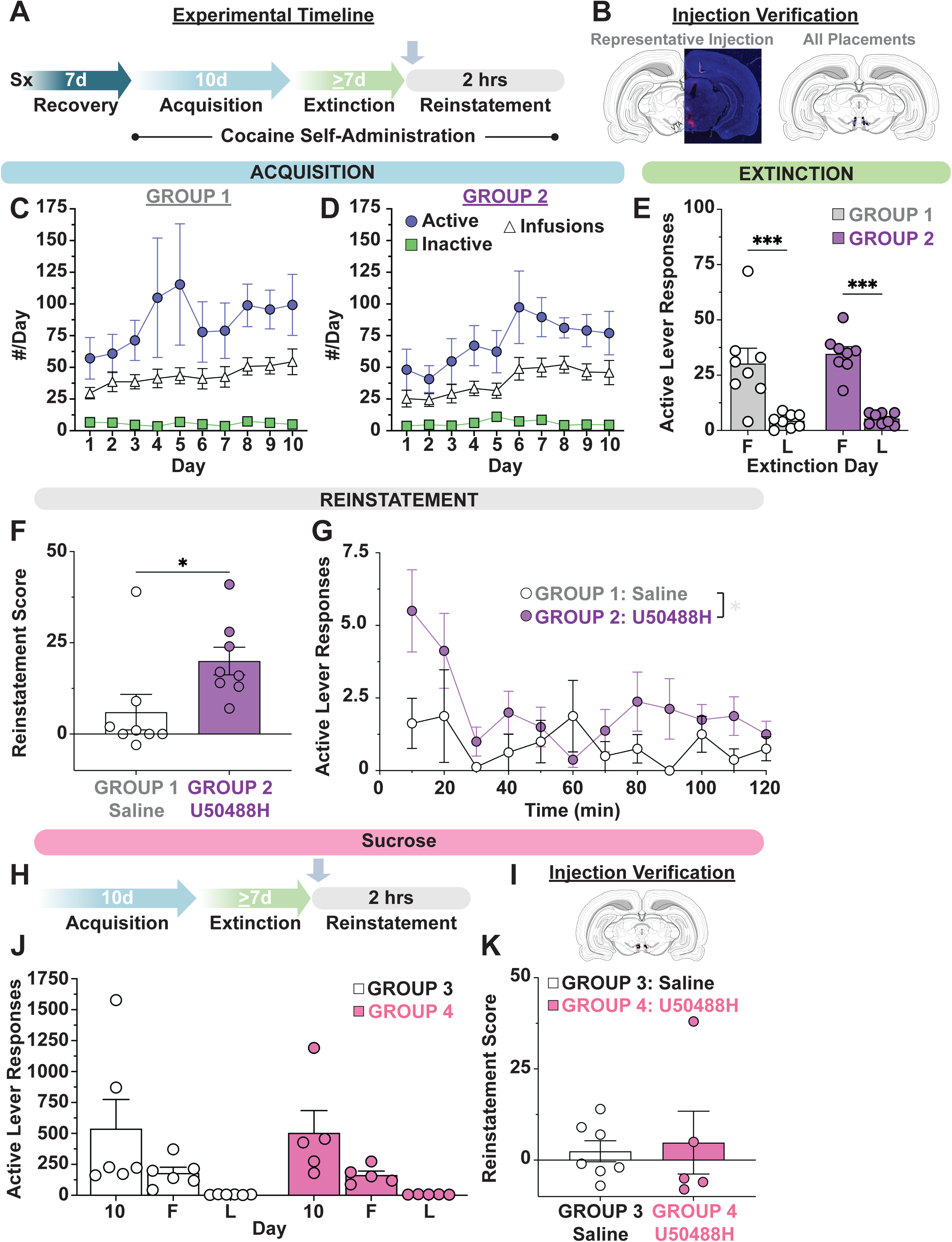
Activation of VTA kORs reinstates cocaine-seeking behavior. (A) Experimental design for cocaine subjects. (B) Cannula placement verification representative image (left) and cannula placements for all animals (right). Purple dots represent U50488H injections, black dots represent control injections. (C, D) Active (blue circles), and inactive (white triangles) lever presses, and cocaine infusions (green squares) for Group 1 (C) and Group 2 (D) animals during 10 days of cocaine self-administration maintenance. There is no difference between groups at this point in the experiment. (E) Active lever presses on the first and last day of extinction training for each group. (n=8 rats/group, 2-way RM ANOVA; main effect of day, p<0.0001). (F) Reinstatement score (calculated as active lever responses on last extinction session subtracted from active lever responses during reinstatement test) of animals given an intra-VTA injection of either U50488H or saline. (n=8 rats/group, Mann-Whitney test, U=8, p=0.01). (G) Active lever responses per ten-minute bin during the 2-hour reinstatement test. (n=8 rats/group, 2-way RM ANOVA, main effect of treatment, p=0.04). (H) Experimental design for sucrose subjects. (I) Cannula placements for sucrose animals. Pink dots represent U50488H injections, black dots represent control injections. (J) Active lever responses for each group on day 10 of maintenance (10), and first (F) and last (L) days of extinction for animals trained to self-administer sucrose pellets. There is no difference between groups at this point in the experiment. (K) Reinstatement scores of animals trained to self-administer sucrose pellets, ten minutes after receiving an intra-VTA injection of either U50488H or saline. * indicates p < 0.05, *** indicates p < 0.0001.

**Figure 8.**
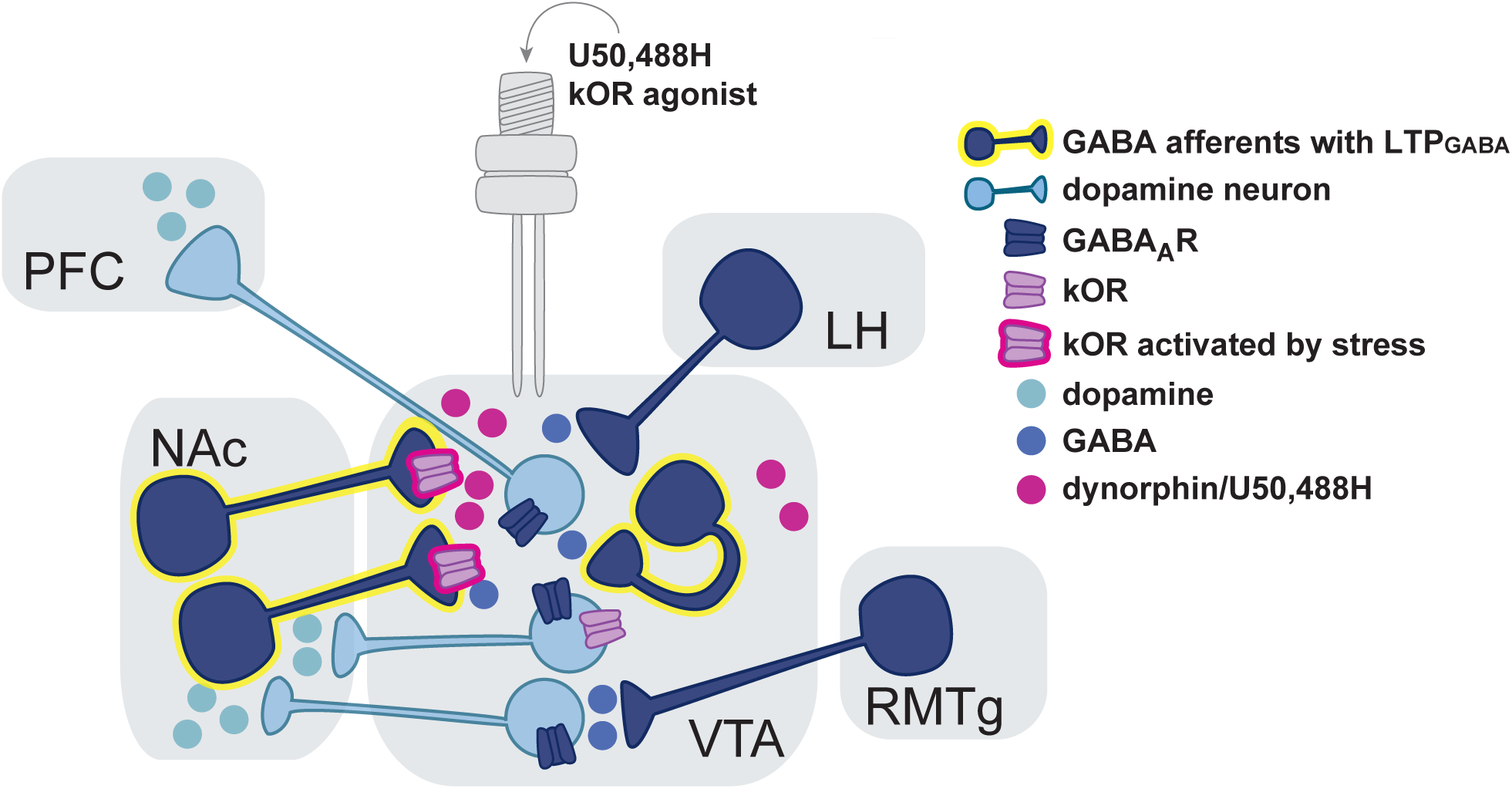
VTA dopamine neurons are controlled by different inhibitory afferents that exhibit distinct synaptic plasticity properties. 1. Stress causes the release of dynorphin into the VTA.
2. Dynorphin binds to kORs on GABAergic terminals from the NAc to block LTP_GABA_. Local VTA GABAergic neurons are similarly affected (Polter et al., 2018).
3. Dopamine neurons express kORs but these do not control LTP_GABA_.
4. Exogenously delivered U50488 mimics stress-induced release of dynorphin, reinstating cocaine-seeking behavior.

To assess the specificity of U50488H on drug-seeking behavior, we also examined its impact on sucrose reinstatement. There was no difference in sucrose self-administration acquisition or extinction between groups (Figure 7J, p=0.89). In addition, intra-VTA microinjection of U50488H did not induce reinstatement of sucrose-seeking behavior (Figure 7K, p=0.53), suggesting that the observed effects may be specific to drug-related rewards.

## DISCUSSION

In earlier work, we were able to correlate acute stress-induced reinstatement of cocaine seeking in rats with the loss of LTPGABA in VTA dopamine neurons (Graziane et al., 2013; Polter et al., 2014, Polter et al., 2017). Our current data show that in mouse as in rat (Morales and Margolis, 2017), the nitric oxide donor SNAP elicits LTP_GABA_, which is also blocked 24 h after acute stress. Moreover (as in the rat), bath application of the kOR inverse agonist nor-BNI to mouse slices rescues LTP_GABA_ 24 h after acute stress. These similarities allowed us to leverage mouse genetics to elucidate the important role in acute stress-induced kOR neuromodulation of GABAergic afferents that project from the NAc to innervate VTA dopamine neurons. We report that 1) synapses from these afferents are normally capable of exhibiting NO-dependent LTP_GABA_; 2) LTP_GABA_ at optogenetically activated NAc-to-VTA synapses is blocked by prior acute swim stress and rescued by *in vitro* treatment with nor-BNI; 3) deletion of kORs selectively from GABAergic NAc neurons rescues LTP_GABA_ 24 h after acute stress; 4) without acute stress, chemogenetically driving pdyn-expressing neurons in the NAc that project to the VTA is sufficient to prevent LTP_GABA_ 24 h later, mimicking what we also observe after acute stress; and 5) that simply activating kORs in the VTA is sufficient to elicit cocaine-seeking in abstinent animals. Our previous work demonstrated interesting properties of kOR receptors in the VTA, i.e. their persistent activation for days following a single exposure to acute stress (Polter et al., 2017). Our current results identify the NAc-to-VTA GABAergic neurons as a likely site of persistent changes in kOR function.

We find that while optogenetically evoked oIPSCs from lateral hypothalamic neurons are not modulated by an NO donor, oIPSCs evoked from NAc neurons exhibit LTP_GABA_. Prior work also supported the idea that LTP_GABA_ is pathway specific, so that SNAP-induced LTP_GABA_ could not be evoked in afferents from RMTg, while synapses from local VTA GABAergic interneurons on neighboring dopamine cells do support LTP_GABA_ (Simmons et al., 2017; Polter et al., 2018). These data suggest that dynorphin released by NAc nerve terminals may not only autoinhibit their own LTP_GABA_, but could spread within the VTA sufficiently to inhibit potentiation of other VTA synapses. Earlier work from our lab found that LTP_GABA_ is not seen at GABA_B_ receptor synapses on VTA dopamine neurons, indicating that NAc afferents targeting these synapses may not serve the same role in stress-induced modulation of behavior (Nugent et al., 2009).

Selective genetic deletion of kORs from dopamine neurons did not prevent acute stress from blocking LTP_GABA_, indicating that the relevant kORs are located elsewhere in the VTA. This finding was surprising in light of previous work on conditioned place aversion (CPA) (Chefer et al., 2013; Ehrich et al., 2015; Abraham et al., 2022). These investigators used a genetic strategy similar to ours to delete kORs from mouse dopamine neurons, and found that CPA was entirely blocked following systemic delivery of the kOR agonist U69593 (Chefer et al., 2013), and in response to a stressor somewhat similar to our 5-minute cold water swim, repeated room temperature swims (Ehrich et al., 2015; Abraham et al., 2022). Moreover, in kOR knockout mice, re-expression of kORs in midbrain dopamine neurons or in the dorsal raphe rescued CPA induced with the kOR agonist (Ehrich et al., 2015; Land et al., 2009). Inactivation of kORs in the dorsal raphe nucleus (Land et al., 2009) or deletion of prodynorphin (Abraham et al., 2022) also blocked stress-induced reinstatement of cocaine CPP. As CPA and CPP are learning paradigms based on the formation of an aversive or appetitive memory, our results may indicate a dissociation between circuitry mediating learned associations between a kOR-mediated aversive event and a specific location and the simpler single, strongly aversive cold water exposure that we used (learning not tested). The circuit elucidated by the previous CPA studies requires involvement of kORs in the dorsal raphe nucleus and kORs present on DAT-expressing neurons.

Recent evidence also indicates that kORs in VTA dopamine neurons are required for the escalation of cocaine consumption (Gordon-Fennell et al., 2023). The strikingly distinct circuit mediating a single cold water swim experience instead requires kORs in NAc-to-VTA GABAergic afferents that regulate the loss of LTP_GABA_, and are correlated with stress-induced cocaine-seeking. Together these results reinforce the idea that kOR activation impacts multiple neural targets related to acute stress and drug-seeking.

Our results show that the NAc could be the *in vivo* dynorphin source relevant to stress induced reinstatement. Consistent with this idea, NAc shell MSNs exhibit LTP at excitatory synapses after cocaine extinction (Ebner et al., 2018) and after aversive experiences including repeated cold water forced swim (Campioni et al., 2009).

Moreover, NAc shell dynorphin-containing neurons are also recruited to drive pain-induced negative affect (Massaly et al., 2019), while the photostimulation of specifically ventral NAc shell elicits robust conditioned and real-time aversive behavior via kOR activation (Al-Hasani et al., 2015). However, this does not rule out the participation of other dynorphin-containing afferents to the VTA that may be recruited during stress.

Projections to the VTA from the dorsal raphe nucleus and the bed nucleus of the stria terminalis have been described to contribute to aversive behaviors such as anxiety and threat generalization (Fellinger et al., 2021); we speculate that projections from these regions may also release dynorphin in the VTA that could trigger stress-mediated loss of LTP and drug-seeking. Interestingly, dynorphin release within the NAc also reduces the activation of VTA-projecting MSNs which release GABA (Tejeda et al., 2017). This is expected to synergize with the loss of LTP_GABA_, further increasing dopamine cell excitability, potentially over several days, and making the individual more susceptible to reinstatement/relapse.

Here we demonstrate for the first time that kOR activation in the VTA is sufficient to induce reinstatement, as intra-VTA microinjection of the kOR agonist U50488H reinstates cocaine seeking. Although we propose that intra-VTA U50488H influences cocaine-seeking behaviors by modulating LTP_GABA_ at synapses on dopamine neurons, there may also be a direct effect on kORs on dopamine neurons themselves. Future investigations will examine whether kOR activation on the NAc terminals is sufficient for stress-induced cocaine reinstatement. We suggest that kORs in the VTA modulate drug reward pathways rather than natural reward systems, since the kOR agonist did not reinstate sucrose-seeking in our experiments. Similarly, VTA-targeted kOR activation does not alter food-related behavioral reinstatement (Sun et al., 2010), although kORs in other brain regions play a role in food-seeking (for review, see Karkhanis et al., 2017). Numerous previous studies have demonstrated that cocaine and sucrose reinstatement often do not align in response to various treatments, both in the VTA and elsewhere (Schmidt et al., 2016; Blacktop & Sorg, 2019). Kappa ORs in several brain regions have been implicated in drug-seeking behaviors. For example,

kOR activation in the NAc core or VTA enhances the aversive properties of nicotine (Pham et al., 2022). Kappa OR inactivation in the basolateral amygdala blocks both U50488H-induced enhancement of nicotine CPP (Smith et al., 2012) and stress-induced reinstatement of nicotine CPP (Nygard et al., 2016). Activation of kORs within the NAc shell inhibits the expression of morphine CPP (Liang et al., 2006), and decreases sucrose self-administration in rats (Massaly et al., 2019), mimicking kOR induced shifts in motivation seen for various drugs of abuse (Liang et al., 2006; Gordon-Fennell et al., 2023). Together these data suggest that kORs in various brain regions may have multiple roles in motivation, depending on whether rewards are present or whether animals are responding for reward-associated cues, and the brain region involved.

Our results are likely specific to stress-induced reinstatement, since systemic kappa antagonist pretreatment does not affect nicotine-primed (Jackson et al., 2012), cue-induced reinstatement of nicotine-seeking (Grella et al., 2014), or context-induced reinstatement of oxycodone seeking (Bossert et al., 2019). Thus, we favor the idea that a unique mechanism underlies stress-induced reinstatement as compared to other forms of reinstatement, which may contribute to the lack of robust therapeutic effects seen with kOR antagonists in clinical trials (for review, see Banks et al., 2020).

## Acknowledgements

The authors would like to thank Dr. Chelsie Brewer for technical and scientific contributions, and Caroline Casper, Junhong Li, and Annie Wolfden for technical assistance. This work was supported by R01 DA 011289 to JAK and TEB, R01 DA 055645 to TEB, Stanford University Dean’s Award (VMD), and Washington State University Alcohol and Drug Abuse Research Program (LGB).

## Author Contributions

All authors had full access to all the data in the study and take responsibility for the integrity of the data and the accuracy of the data analysis. Study concept and design: J.A.K. and T.E.B. Acquisition of data: V.M.D., L.G.B. and J.S. Analysis and interpretation of data: V.M.D., L.G.B., J.S., J.A.K. and T.E.B. Original drafting of the manuscript: V.M.D. and L.G.B. Critical revision of the manuscript for important intellectual content: J.A.K., T.E.B., V.M.D., L.G.B. Statistical analysis: V.M.D. and L.G.B. Funding acquisition: J.A.K. and T.E.B. Administrative, technical, and material support: J.S. and A.T. Study supervision: J.A.K. and T.E.B.

## Supplemental Figures

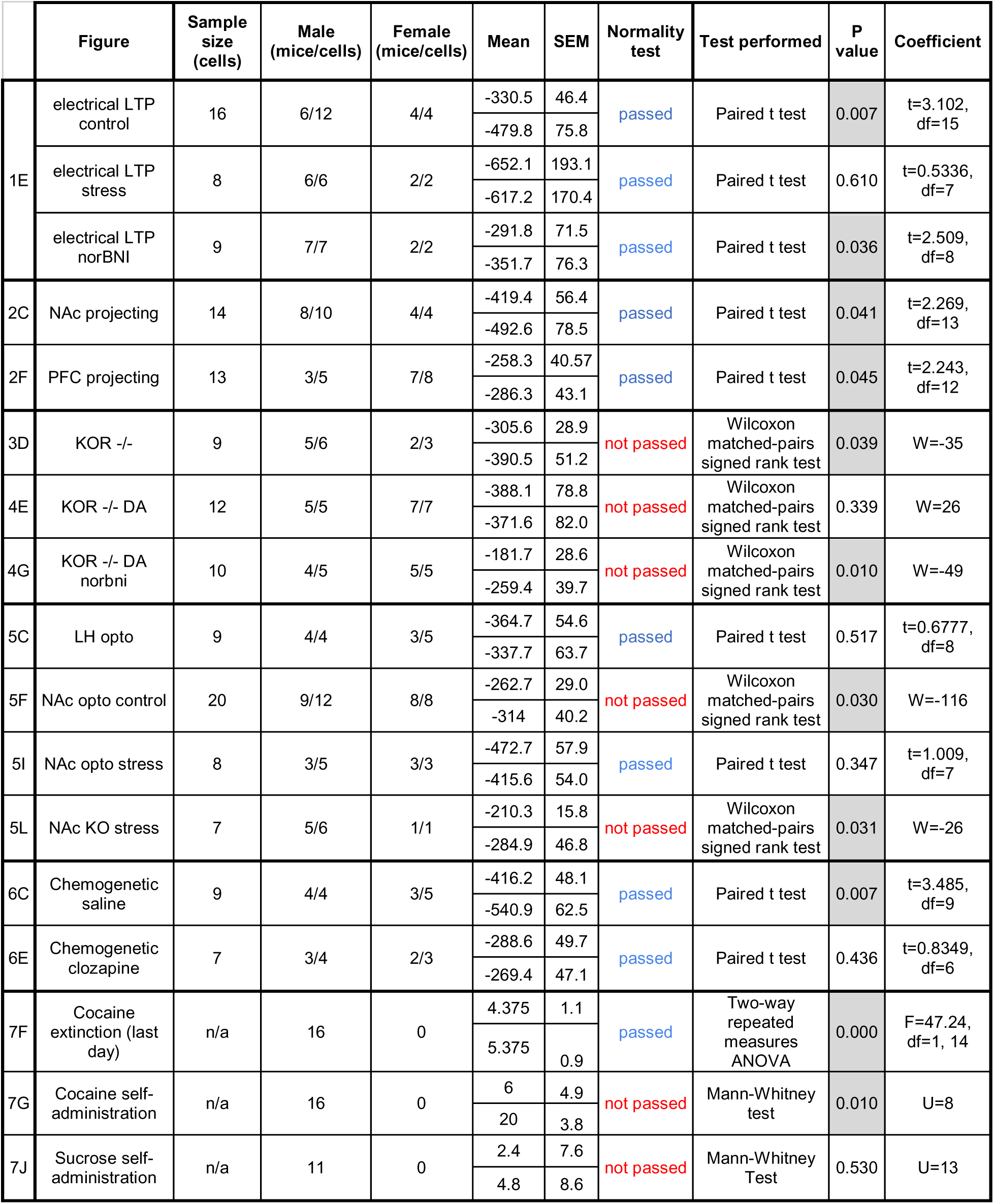

**Supplemental Figure 1.**
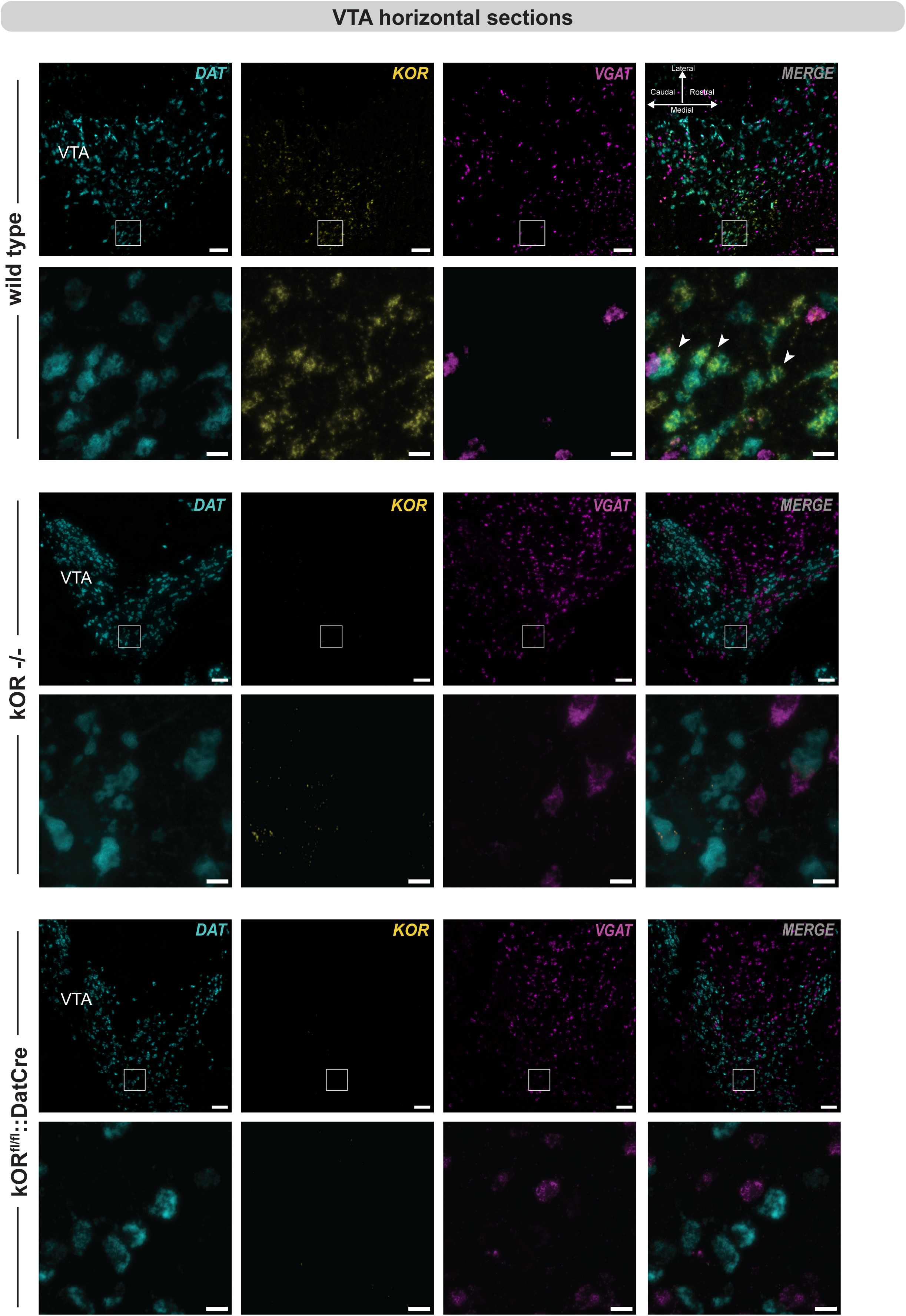
Validation of mouse lines and strategies used for assessing kOR relevant location via *in situ* hybridization in the VTA. Representative confocal images of fluorescent *in situ* hybridization on horizontal slices through the VTA of wild type mice expressing kOR in Dat+ neurons, kOR ^-/-^ mice that lack kOR in every cell and brain region that naturally expresses it, and Dat-cre::Oprkfl/fl mice that lack kOR only in Dat+ neurons. The presence of different transcripts is indicated as follows: Dat (cyan), Kor (yellow), Vgat (magenta), and merge (overlay of all 3 signals). Scale bar: low magnification = 200 μm, high magnification = 30 μm.

**Supplemental Figure 2.**
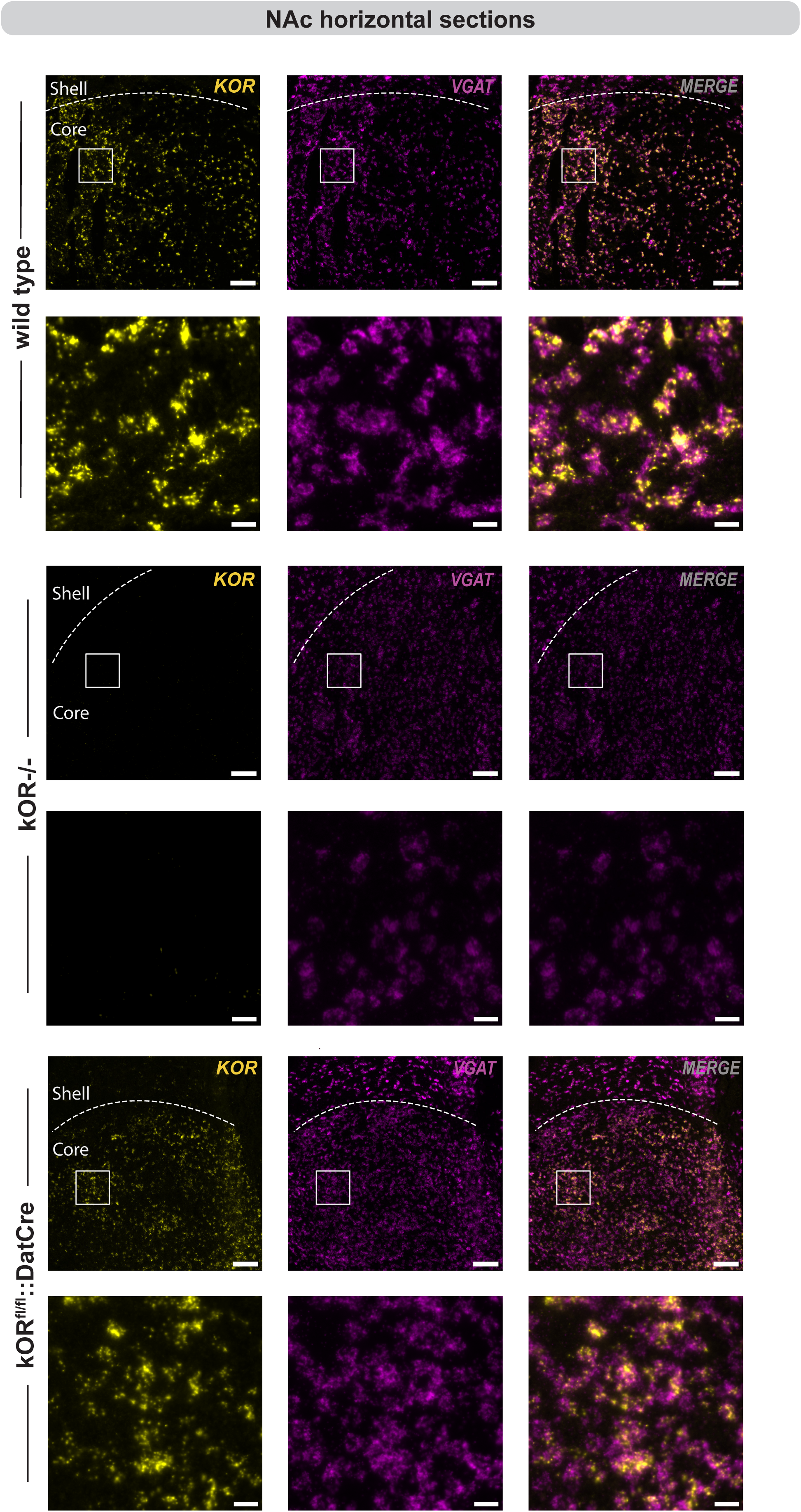
Validation of mouselines and strategies used for assessing kOR relevant location via *in situ* hybridization in the NAc. Representative confocal images of fluorescent *in situ* hybridization in horizontal slices through the NAc of (A) wild type mice expressing kOR in Vgat+ neurons, (B) kOR ^-/-^ mice that lack kOR in every cell and brain region that naturally expresses it, (C) Dat-cre::Oprkfl/fl mice that lack kOR only in Dat+ but not in Vgat+ neurons. The presence of different transcripts is indicated as follows: Dat (cyan), Kor (yellow), Vgat (magenta), and merge (overlay of all 3 signals). Scale bar: low magnification =200 μm, high magnification = 30 μm.

